# Sex-biased dispersal drives mito-nuclear discordance in simulated populations

**DOI:** 10.1101/2025.05.19.654820

**Authors:** Francesco Giannelli, Joan Ferrer Obiol, Emiliano Trucchi

## Abstract

Reconstructing the evolutionary dynamics of natural populations requires an understanding of the geographical distribution of nuclear and mitochondrial genetic diversity. The analysis of these two genetic markers frequently discloses discordant patterns (mito-nuclear discordance) that can arise simply as a consequence of their different effective population sizes (*N*_e_). Species-specific sex-biased dispersal may contribute to the mito-nuclear discordance observed in natural populations. However, the relative contribution of genetic drift versus sex-biased dispersal in driving mito-nuclear discordance remains insufficiently evaluated. Here, we use forward genetic simulations to address this knowledge gap. Our findings support the baseline level of mito-nuclear discordance arising from distinct genomic *N*_e_, but show that this inherent discordance is magnified by sex-biased dispersal patterns. We demonstrate that female-biased dispersal leads to a marked spatial mismatch between mitochondrial and nuclear diversity across the simulated populations, thereby reducing the spatial concordance between mitochondrial and nuclear markers. Conversely, male dispersal patterns appear to increase, although to a reduced degree, the intrinsic level of discordance between nuclear and mitochondrial geographical marker distribution. Our results highlight the importance of integrating the intrinsic characteristics of nuclear and mitochondrial genomes and the impact of sex-biased dispersal for accurately interpreting patterns of genetic diversity and reconstructing evolutionary histories.

## Introduction

### MITO-nuclear discordance impacts Population Genetic Inference

Mitochondrial and nuclear markers have a long history of being employed to assess spatial patterns of genetic variation within and among populations. Despite being widely used for phylogenetic and phylogeographic analyses, these markers differ in their basic features. The mitochondrial genome, due to its maternal inheritance and its haploidy, has an effective population size four times smaller and reduced gene flow compared to the nuclear genome (Palumbi, Cipriano, and Hare 2001; Hudson and Turelli 2003; Zink and Barrowclough 2008). Furthermore, the mitogenome does not recombine and has a higher evolutionary rate due to the lack of DNA repair mechanisms, resulting in a faster accumulation of mutations (Lynch, Koskella, and Schaack 2006; Galtier et al. 2009). This implies that demographic processes may affect mitochondrial and nuclear markers differently, which can lead to a lack of consistency in the geographical distribution of genetic diversity. This inconsistency has been acknowledged in several studies as mito-nuclear discordance and has been reported with increasing frequency as long as the availability of nuclear genome data in phylogeographical studies increased (Toews and Brelsford 2012). Different mechanisms have been proposed to explain mito-nuclear discordance: (*i*) demographic processes that enhance genetic drift (Petit and Excoffier 2009); (*ii)* introgression (Cahill et al. 2013; Morales et al. 2015); (*iii*) incomplete lineage sorting (Maddison 1997; Suh, Smeds, and Ellegren 2015; de Jong et al. 2023); (*iv*) selective processes (Morales et al. 2015) and (*v*) sex-biased asymmetries (Toews and Brelsford 2012). As of today, despite the widespread reporting and description of mito-nuclear discordance in many natural populations and taxa, the use of a single genetic marker, either nuclear or mitochondrial, remains common when describing genetic diversity and population structure. This practice can result in a partial or incorrect depiction of the geographic dynamics of populations.

### Sex-biased dispersal patterns in natural populations

Among sex-biased asymmetries, one phenomenon that has often been accounted as a possible explanation for the occurrence of mito-nuclear discordance is sex-biased dispersal (Toews and Brelsford 2012; Dessi et al. 2022; Dupont et al. 2022). This term refers to the presence of different patterns of dispersion between individuals of different sexes within a population (Li and Kokko 2019). According to literature, sex-biased dispersal appears to be a consequence of specific animal behaviours like (i) mate competition, (ii) resource competition, (iii) inbreeding avoidance (iv) kin competition, (v) mortality costs, (vi) mating systems, and (vii) environmental and/or demographic stochasticity (Greenwood 1980; Trochet et al. 2016; Li and Kokko 2019). Two contrasting patterns of sex-biased dispersal are observable in nature: male or female-biased dispersal, where males or females exhibit greater dispersal rates, respectively. In each vertebrate taxon, there are documented occurrences of either behaviour, but a general trend is often recognisable: male-biased dispersal is more often reported in mammals (Greenwood 1980; Stephen Dobson 1982; Trochet et al. 2016), while in contrast, birds show the opposite pattern (Greenwood 1980; Mabry et al. 2013; Peters et al. 2014; Trochet et al. 2016; Palacios et al. 2023). In fishes (Hutchings and Gerber 2002; Taylor et al. 2003; Anseeuw et al. 2009; Cano, Mäkinen, and Merilä 2008) and reptiles (OLSSON and SHINE 2003; Keogh, Webb, and Shine 2006; Dubey et al. 2008; Ujvari, Dowton, and Madsen 2008) both patterns are present to the same extent.

### Aims of this study

In 2012, Toews and Brelsford identified the study of the drivers of mito-nuclear discordance as a critical focus. Yet, understanding the origin of such a pattern has not received adequate attention. Furthermore, the impact of the potential drivers of discordance has not been thoroughly estimated or studied, either empirically or through simulation studies. In this study, we investigated whether differences in effective population size alone could explain mito-nuclear discordance or if incorporating sex-biased dispersal into the model is necessary to fully capture the observed patterns of genetic diversity. Our goal is to understand how the two markers are capable of describing the structure of a population and how they correlate with each other. Our focus is on sex-biased dispersal as an explanation for the occurrence of mito-nuclear discordance in natural populations (Toews and Brelsford 2012; Dessi et al. 2022; Dupont et al. 2022). We hypothesise that both male-biased and female-biased dispersal may impact the correlation between nuclear and mitochondrial markers, potentially resulting in a decreased correlation between the two. Specifically, we hypothesise that because mitochondrial DNA is only passed from mothers, male-biased dispersal will not directly affect the distribution of mitochondrial markers but will influence only nuclear markers. As a result, we expect this dispersal pattern to have a minor overall effect on the discordance between nuclear and mitochondrial genomes. In contrast, when females disperse, the movement of individuals results in the redistribution of both mitochondrial and nuclear markers across populations, leading to a more pronounced disruption in their correlation.d

## Materials and Methods

### Spatially explicit forward simulations design

To test our hypotheses, we created a two-dimensional forward simulation scenario using SLiM v4.0.1 (Haller and Messer 2023). To design a more ecologically realistic model, we based our simulations on non-Wright-Fisher assumptions (Haller and Messer 2019), as suggested for spatial models. This allowed us to simulate a population with overlapping generations, age-based mortality and population size (N) as an emergent parameter. As we wanted N to be generally stable around a pre-determined size and population density to be constant, we set up a carrying capacity (K) value of 20 ‘000, which translates into a stable N value and population density, resulting from the equation N = K / Competition (Dieckmann and Doebeli 1999; Doebeli and Dieckmann 2003). Competition directly affects individual fitness, which in this context depends solely on the surrounding population density rather than genetic mutations. As density increases, competition becomes more intense, reducing local individual fitness. We enabled the presence of different sexes in the simulations, with the sex of every newborn individual being determined by the SLiM default option to maintain the sex ratio around 0.5.

Each simulated individual carries a genome composed of ten nuclear fragments of 160 ‘000 bp and one mitochondrial portion of 16 ‘000 bp for a total of 1 ‘616 ‘000 bp. This results in the nuclear portion of the genome being one thousand times larger than the mitochondrial one. We set the nuclear genome’s mutation rate at the SLiM default of 1×10^−8^, while the mitochondrial genome’s mutation rate was set 10 times faster than the nuclear one, at 1×10^−7^, following Allio et al. (2017), while the recombination rate has been set as 1e-8 within the nuclear genome and as 0 within the mitochondrial genome. Our simulation model does not involve selection dynamics, so all mutations are generated as completely neutral. It is also important to mention that the nuclear and mitochondrial components of the genomes are completely independent of each other as well as each one of the ten nuclear portions. We also implemented the sole maternal inheritance of the mitochondrial genome in the simulation. We enforced strict maternal inheritance by assigning mitochondrial genetic material to chromosome 1 in SLiM. For chromosome 2—representing the paternal contribution—we systematically removed any mitochondrial mutations, ensuring that only the maternally derived mitochondrial genome was transmitted.

To explore the spatial distribution of the different markers, we created a homogenous two-dimensional area of 1 × 1 in which individuals disperse in their first year of life starting from their mother’s spatial position within a radius distance of 0.004. In this model, individuals reproduce according to their proximity, ensuring that breeding only occurs when a potential mating partner is within the range of interaction of 0.008.

### No sex-biased dispersal model

In the first simulated scenario, we randomly distributed 20’000 individuals over the two-dimensional area (Figure 1. A). In this model, the dispersal rate is the same for both sexes, a 0.004 radius from their mother’s spatial position, happening only in the individual’s first year of life. This model has been created to understand the expected degree of mito-nuclear discordance due to the sole intrinsic characteristics (i.e., effective population size) of the two genomes and to serve as a reference for the other models. We ran this model for 100’000 generations, performing 10 independent replicates.

**Figure 1.**
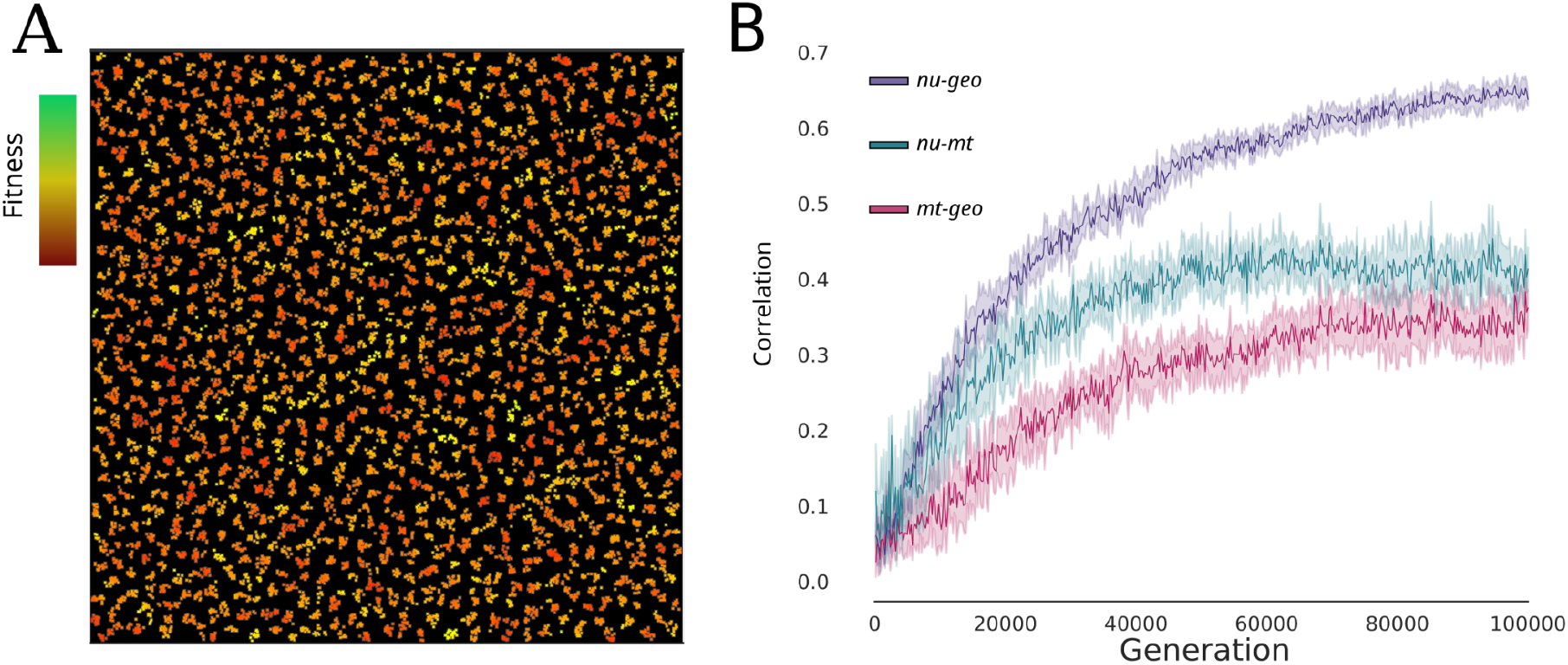
**A –** Graphical representation of the simulated individuals in the bi-dimensional area. Individuals are coloured according to relative fitness, which in this model, is derived from the local population density. **B) –** Trajectory of the Pearson correlation values between I) *nu-geo* (in purple), II) *nu-mt* (in green) and III) *mt-geo* (in pink) from generation 0 to generation 100’000.

### Sex-biased dispersal model

To explore the impact of sex-based dispersal on the geographical distribution of mitochondrial and nuclear genetic diversity, we created six different models, three of which featured a female-biased dispersal pattern in which females, in their first year of life, dispersed twice, three times, four times and five times as much as male individuals (respectively 0.008, 0.012, 0.016 and 0.0020). In the other three models, we simulated the same pattern but reversed in favour of the males dispersing more than the females (Table 1). Again, each model ran for 100’000 generations, with 10 replicas each.

**Table 1.** Female and male dispersal rates in each simulation model we tested. Non sex-biased dispersal model is indicated as NDB. FBD stands for female-biased dispersal, and MBD for male-biased dispersal.

|  | NBD | FBD | FBD2 | FBD3 | FBD4 | MBD | MBD2 | MBD3 | MBD4 |
| --- | --- | --- | --- | --- | --- | --- | --- | --- | --- |
| Female dispersal rate | 0.004 | 0.008 | 0.012 | 0.016 | 0.0020 | 0.004 | 0.004 | 0.004 | 0.004 |
| Male dispersal rate | 0.004 | 0.004 | 0.004 | 0.004 | 0.004 | 0.008 | 0.012 | 0.016 | 0.0020 |

### Sampling and statistics calculation

To track the evolution through time of the correlation between nuclear and mitochondrial genetic diversity and geometric distance, we set up a systematic sampling scheme: every 200 generations we randomly sample 100 individuals from the population. We then calculated the number of unshared nuclear and mitochondrial mutations between every possible couple of individuals in the sample and measured the geometric distance between them. At the end of every simulation, we calculate the Pearson correlation (Pearson and Lee 1903) between:

- Nuclear genetic diversity and geometric distance (*nu-geo*)
- Mitochondrial genetic diversity and geometric distance (*mt-geo*)
- Nuclear genetic diversity and mitochondrial genetic diversity (*nu-mt*)

As stated before, each model has been replicated 10 times using a different starting seed number. Seed numbers set the random parameters governing the simulation events, ensuring the robustness and reliability of our analyses across multiple replicas.

## Results

### No sex-biased dispersal scenario

Our simulations incorporating the NBD pattern demonstrate the progressive development of an isolation-by-distance (IBD) pattern in both nuclear and mitochondrial genomes (Figure 1. B). Importantly, even in the absence of any sex-biased dispersal pattern, the *mt-geo* correlation is always lower than the *nu-geo* correlation. The same applies to *nu-mt* which stands between the other two correlations (Figure 1. B). This result already confirms that a certain degree of discordance between mitochondrial and nuclear genomes is expected even without sex-biased dispersal. As theoretically predicted, this is the result of the intrinsic characteristics of the mitochondrial genome: smaller genome size, smaller population size and lack of recombination, which are reflected in a generally lower number of unshared mutations between individuals (Supplementary Figure 1 and 2). Such a reduced number of mutations appears to be a limiting factor when describing a natural population’s structure and creates an inconsistency with the nuclear genome, which, instead, holds more information and is, therefore, a finer descriptor of the geometric distances between individuals.

**Figure 2.**
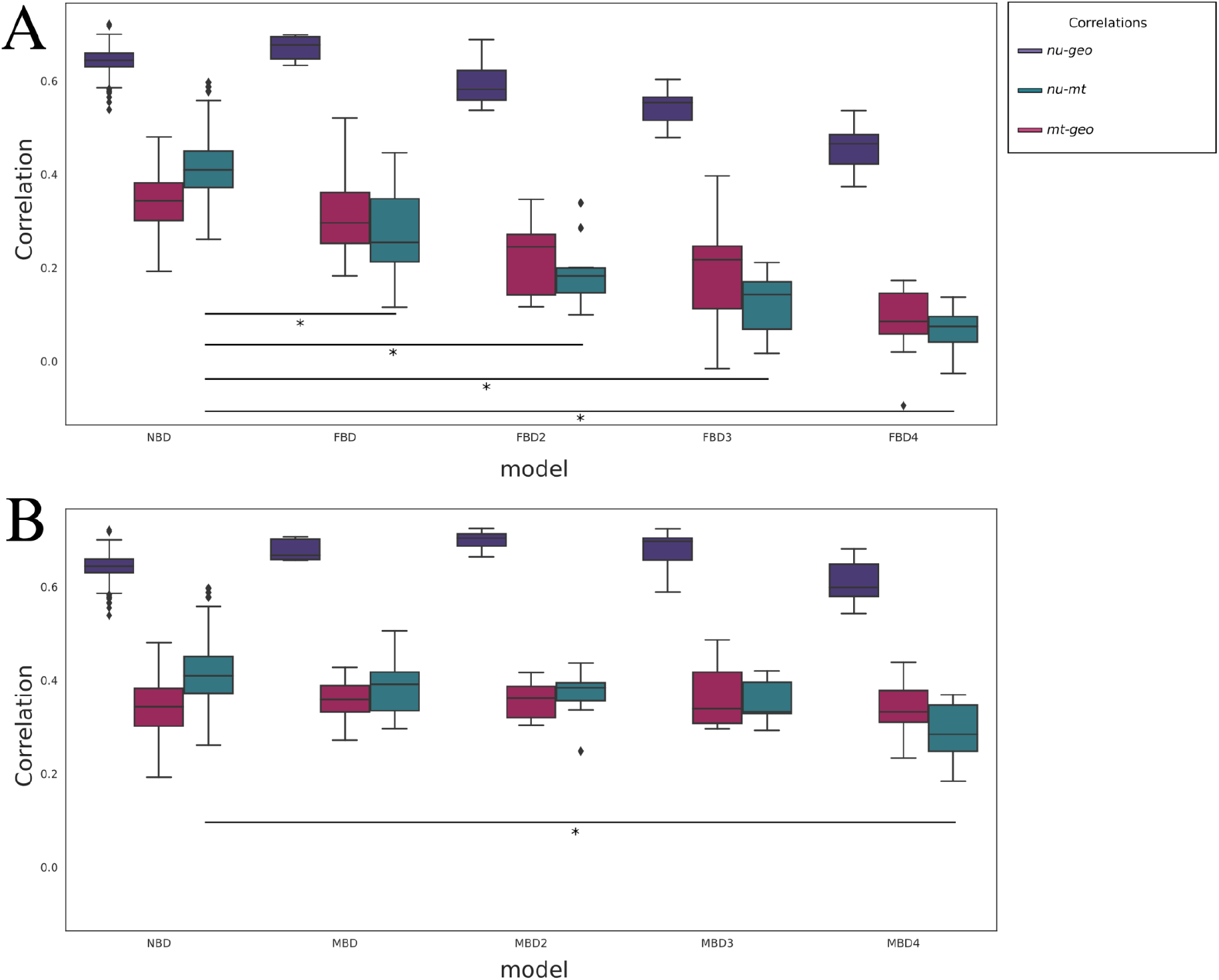
**A –** Pearson correlation values between the different female-biased dispersal patterns calculated on the results of the last 10’000 generations (from generation 90’000 to generation 100’000). On the left, model NBD is serving as a reference, then, moving from left to right, we have the results from FBD, FBD2, FBD3 and FBD4 models. **B** – Pearson correlation values between the different male-biased dispersal patterns in the last 10’000 generations. On the left, model NBD serves as a reference, then moving from left to right, we have the results from MBD, MBD2, MBD3, MBD4 models.

### Sex-biased dispersal

#### Female-biased dispersal

The *nu-mt* correlation reaches its lowest levels in the models featuring female-biased dispersal. In these models, the *nu-mt* correlation is inversely proportional to the female dispersal rate (Figure 2.A). The difference in the value distribution is significantly different for all the models compared to the non-sex-biased dispersal model. This observed reduction in the *nu-mt* correlation is primarily driven by mitochondrial genetic diversity failing to accurately reflect geometric distances (Supplementary Figure 2). This outcome aligns with theoretical expectations, as the increased dispersal rate of females interferes with establishing a strong isolation-by-distance (IBD) pattern for the mitochondrial genome. This pattern arises due to the absence of recombination in the mitochondrial genome, which limits the retention of newly arising mutations over time. As a result, the mitochondrial population structure is defined by a very small number of mitohaplotypes, differentiated by only a few mutations (Supplementary Figure 2).

We also observe a decline in the *nu-geo* correlation, which is directly correlated to the female dispersal rate. This pattern is likely driven by the overall increase in individual dispersal caused by female movement. Since offspring are generated starting from their mother’s position, higher female dispersal leads to a greater mixing of genetic diversity across spatial scales, thereby weakening the correlation between nuclear diversity and geometric distance.

#### Male-biased dispersal

In scenarios characterised by male-biased dispersal (Figure 2.B), a significant reduction in the *nu-mt* correlation is observed only under conditions where males disperse at a rate five times higher than females (MBD4). This suggests that male-biased movement alone does not substantially impact the relationship between nuclear and mitochondrial diversity unless the disparity in dispersal rates is extreme. Furthermore, the *mt-geo* correlation remains largely unaffected across all male-biased dispersal scenarios, as mitochondrial diversity is inherited maternally and, therefore, remains predominantly shaped by female movement rather than male dispersal patterns.

## Discussion

This study highlights how nuclear and mitochondrial genomes have intrinsically different descriptive powers concerning population structure. Our findings demonstrate that a certain amount of discordance between the two markers is always present. However, a correlation between sex-biased dispersal and mito-nuclear discordance exists and is much more pronounced in models in which the female is the sex with a higher dispersal rate. This leads us to predict that mito-nuclear discordance may be particularly frequent in taxa in which female-biased dispersal is predominant like birds (Toews and Brelsford 2012; Palacios et al. 2023; Peters et al. 2014) where females (ZW) are the heterogametic sex and, as hybrid offspring, are expected to exhibit reduced fitness according to Haldane’s rule (Orr 1997).

Our results show that the mitochondrial genome always has a lower descriptive power compared to the nuclear genome, due to its intrinsic characteristics. This is also supported by our results where, in the case of sex-biased dispersal patterns in a homogeneous and stable environment, mito-nuclear discordance is mainly driven by the mitochondrial genome being a poorer descriptor of population’s structure. Nevertheless, this does not imply that this marker should not be taken into account, but rather that, if correctly interpreted, it can offer crucial insights into ongoing evolutionary and demographic processes that complement the information provided by the nuclear genome, and vice versa (Morales et al. 2015; DeRaad et al. 2023; Chiocchio et al. 2024). As other researchers have pointed out in previous papers (Folt et al. 2019), it is fundamental to use both markers separately when describing population structures and contextualise the results according to the species’ behavioural characteristics, the evolutionary history of the populations, and the environmental parameters that may have influenced the distribution of genetic diversity within the population under study.

In this study, we intentionally refrained from exploring the potential causes of sex-biased dispersal, despite their recognised importance in shaping population dynamics (Li and Kokko 2019). While understanding these underlying drivers is essential for a comprehensive interpretation of dispersal patterns, their investigation falls beyond the scope of our work. Instead, we focus solely on the outcomes of these dynamics. More broadly, this study represents the first attempt to use simulation-based approaches to examine the relationship between mito-nuclear discordance and one of its proposed underlying mechanisms.

### Model limitations

Like every other simulated model, the scenarios we created have intrinsic limitations. In particular, regarding the dispersal parameter, we tried to use realistic dispersal distances that could produce IBD patterns in our models. The interactions between individuals have been set to avoid unrealistic local population densities and reproduction distances. Still, we are conscious that it is impossible to have a clear estimation of the intensities of the interactions between individuals and the maximum distance at which these interactions continue to take effect.

Other than that, we can see that, in female-biased dispersal scenarios, the *nu-geo* correlation is inversely proportional to the females’ dispersal rate. This is because every newborn is generated starting from the female’s spatial position. This results in an overall increased dispersion of the nuclear marker in the case of female-biased dispersal, which, over a certain threshold, decreases the *nu-geo* correlation.

### Future perspectives

In our work, we explored a reduced set of parameters to minimise the emergent effects of combining different dynamics, which might otherwise confound the interpretation of our results. We are aware that natural populations’ actual dynamics are much more complex, involving intricate interactions between biological and non-biological components. One interesting dynamic to include in future exploratory models is changes in population size, which have been described as the potential causative agent of mito-nuclear discordance (Larmuseau et al. 2010). Additionally, implementing different mating systems and migratory dynamics (seasonal and non-seasonal) in our models would help us understand how these dynamics may interact with differential dispersal patterns. Our simulations were conducted under the assumption of a homogeneous and stable environment. However, we recognise that over long timescales environmental conditions can fluctuate, and such dynamics have been proposed as contributing factors to mito‐nuclear discordance (de Jong et al. 2023). Finally, a valuable way to test the robustness of our model would be to apply it to real-world data, incorporating accurate information on the movement of individuals, population density, reproductive systems, and migration patterns. This approach would allow us to validate our findings and refine our model to reflect the complexities of natural populations better.

## Acknowledgements

We warmly thank Josephine R. Paris, Marco Gargano, Francesco Maroso, Giorgio Bertorelle, Sebastiano Fava, Lorena Ancona, Flavia A.N. Fernandes, and Samuele Broccardo for their invaluable support throughout the development of this work. Their contributions, advice, and stimulating discussions have been essential to the completion of this study. Simulations were performed on the HPC clusters at the Department of Life and Environmental Sciences (“HappyComputing@DiSVA”), Marche Polytechnic University, Italy. We also acknowledge the use of AI-assisted tools (ChatGPT by OpenAI) to improve the clarity and language of the manuscript.

## Data availability

The scripts are available at: https://github.com/giannef/Simulations-mito-nuclear-discordance

## Contributions

E.T. conceived and supervised the study. F.G. designed the models, conducted the analyses, and wrote the manuscript. J.F.O. contributed to data analysis and interpretation.

