## Supplementary materials for "Sex-biased dispersal drives mito-nuclear discordance in simulated populations"

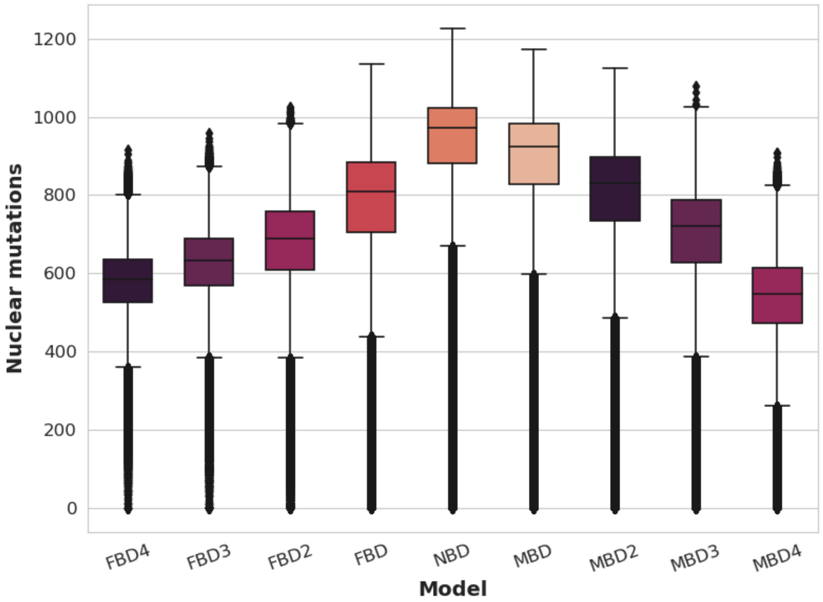


**Supplementary Figure 1.** Distribution of pairwise unshared nuclear mutations across tested models. Boxplots represent the number of nuclear mutations not shared between pairs of individuals for each demographic model tested (X-axis). The Y-axis indicates the count of unshared mutations per pair.

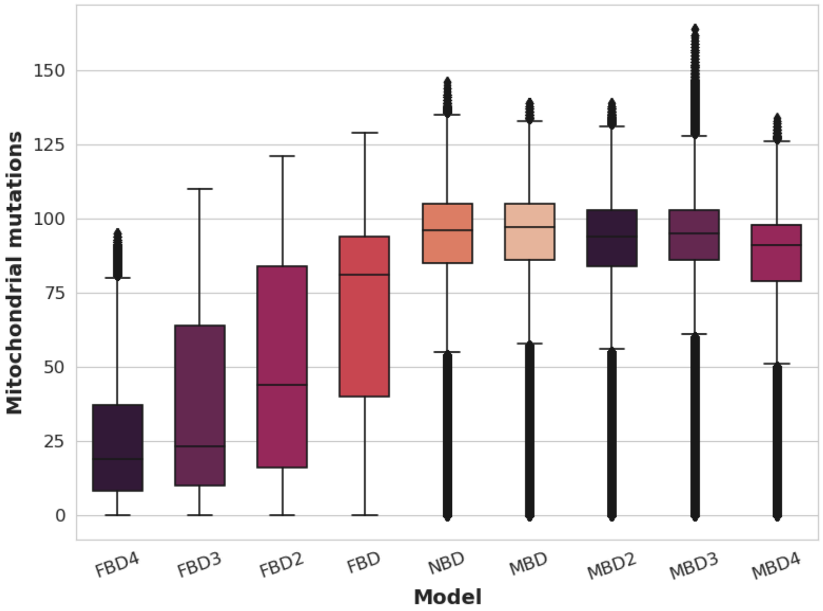


**Supplementary Figure 2.** Distribution of pairwise unshared mitochondrial mutations across tested models. Boxplots represent the number of mitochondrial mutations not shared between pairs of individuals for each demographic model tested (X-axis). The Y-axis indicates the count of unshared mutations per pair.
